# *In situ* Membrane Protein Expression by Efficient Recruitment of mRNA to the Membranes of Synthetic Cells

**DOI:** 10.1101/2024.11.23.624830

**Authors:** Hang Fu, Lijuan Ma, Chunhua Xu, Jinghua Li, Yunhai Sun, Yuanxiao Tao, Hao Wang, Shuxin Hu, Meifang Fu, Hai Zheng, Chenli Liu, Fangfu Ye, Yan Qiao, Ming Li, Ying Lu

## Abstract

The synthesis of artificial cells is crucial for understanding the origins of life. In synthetic cells, however, the absence of membrane-bound organelles and auxiliary proteins severely limits the efficient expression and precise localization of membrane proteins using cell-free expression systems. Here we introduce a robust method that significantly enhances membrane protein synthesis by recruiting mRNA to phospholipid membranes. This is achieved through the use of cholesterol-modified, single-stranded DNA that anchors to the membrane and pairs with the mRNA’s untranslated region. This strategic placement facilitates the assembly of protein expression machinery, promoting direct co-translational folding at the membrane. Our approach not only ensures correct protein topology and functionality but also demonstrates broad applicability for the *in situ* expression of various membrane proteins. It effectively addresses the challenges of membrane protein localization and assembly in synthetic cells, showcasing its versatility for synthesizing a wide array of membrane proteins.

## Main

Synthetic cells, designed to mimic the functions and behaviors of natural cells, represent a pivotal area of modern synthetic biology with promising applications across various fields. Advancements in molecular biology and synthetic biology have propelled artificial cell technology to the forefront of biological research. Like their natural counterparts, synthetic cells are enclosed by membranes equipped with membrane proteins that play essential roles in cellular functions, including signaling, material transport, energy production, and membrane structuring^1^. The synthesis and precise localization of these proteins are essential in synthetic cells, where they enable synthetic cells to emulate fundamental biological functions, thus enhancing their integration and functionality within biological systems^2–4^.

The synthesis and localization of membrane proteins in synthetic cells, particularly when using cell-free systems, pose significant challenges. The inherently hydrophobic nature of these proteins often leads to misfolding and toxic aggregation in aqueous environments^5,6^. In contrast, natural cells employ co-translational pathways to mitigate these issues. Notably, the signal recognition particle (SRP) pathway facilitates direct insertion of α-helical membrane proteins into membranes, substantially reducing the risk of cytoplasmic aggregation^7–12^. However, replicating these complex pathways in synthetic systems is non-trivial. The insertion of hydrophobic amino acids in a random coil conformation into the membrane requires overcoming a substantial thermodynamic energy barrier^13^. The lipid head groups at the lipid bilayer surface promote α-helix formation in peptide chains, lowering the energy barrier to membrane integration^6,14–16^. Previous studies have shown that reducing the size of synthetic cells, mimicking chaperones, and utilizing specialized ribosomes can effectively increase the expression of membrane proteins in membranes^17–19^.

In synthetic cells, where ribosomes must be shared for the expression of various proteins and cell sizes cannot be sufficiently small, it is essential to implement methods that directly enhance the expression of membrane proteins. Researchers have long observed the non-uniform distribution of mRNA within cells^20^. The localization of mRNAs to subcellular compartments provides a mechanism for spatial regulation of gene expression^21,22^. It has been observed that mRNA coding for membrane proteins in eukaryotes tends to be co-located with the endoplasmic reticulum (ER)^23–26^. Analysis of this mechanism revealed that the localization of a subset of mRNA is independent of ribosomes^27^. In prokaryotes, certain mRNA uses their coding region sequences and structures to interact with the SecYEG translocon. This interaction allows the mRNA to be targeted to the membrane independently of the SRP and ribosomes. This mechanism is regarded as a bacterial strategy for synthesizing membrane proteins in response to stressful environmental conditions^28,29^.

In this work, we developed a strategy applicable for efficient synthesis and localization of membrane proteins in synthetic cells based on modifications to the mRNA’s untranslated regions (UTRs). Our approach adds a specially engineered sequence, termed “AnchorTail” (Supplementary Table 1), to the 3’ UTR. This sequence is designed to bind to a cholesterol-modified ssDNA, effectively recruiting the mRNA to the membrane and significantly increasing the probability of co-translational membrane insertion. This method does not require alterations to the protein sequence nor customized translation machinery. We synthesized a few membrane proteins with different number of transmembrane regions that could form both monomers and dimers. The synthesized proteins demonstrated enhanced localization and functionality, making this technique a significant advancement for the construction of synthetic cells capable of sophisticated interactions with their environments.

## Results

### Recruitment of mRNA for membrane protein expression to the membrane via cholesterol-tagged complementary single-stranded DNA

As a proof of concept, we evaluated the feasibility and efficiency of our method on supported lipid bilayers (SLBs) using the ATP Synthase F_0_ subunit a from *E. coli* (gene name: atpB) as a model system. The F_0_ subunit a contains five transmembrane helices, and proper binding to the membrane results in the N- and C-termini being exposed on opposite sides (Supplementary Figure 1), which is a characteristic that can be used to assess the orientation and localization of the membrane proteins.

We created a DNA construct that encodes H6-atpB-Flag-AnchorTail (Fig.1a), which includes the F_0_ subunit a protein tagged with His×6 at the N-terminal and FLAG at the C-terminal. The AnchorTail, located in the 3’ UTR, consists of a 60-base pairs sequence designed to make the transcribed mRNA hybridize with the cholesterol-labeled ssDNA (chol-ssDNA). We synthesized and purified the H6-atpB-Flag-AnchorTail mRNA in vitro, allowing it to anneal with the cholesterol-labeled ssDNA and a Cy3-labeled ssDNA (Cy3-ssDNA) in a 1:1:1 ratio, resulting in the formation of membrane-bound mRNA that we named chol-ssDNA/mRNA (Supplementary Figure 2, Fig.1b). The Cy3-labeled ssDNA was used to visualize the mRNA localization. As a control, we used an identical sequence without the cholesterol label, maintaining the same nucleic acid structure as the experimental group but lacking the membrane binding capacity. Use of the bulk PURE system with our construct enabled the synthesis of H6-atpB-Flag from mRNA, confirmed by Western blot analysis (Supplementary Figure 3). The results demonstrated the ability of our nucleic acid construct to be used as a translation template.

**Fig. 1.**
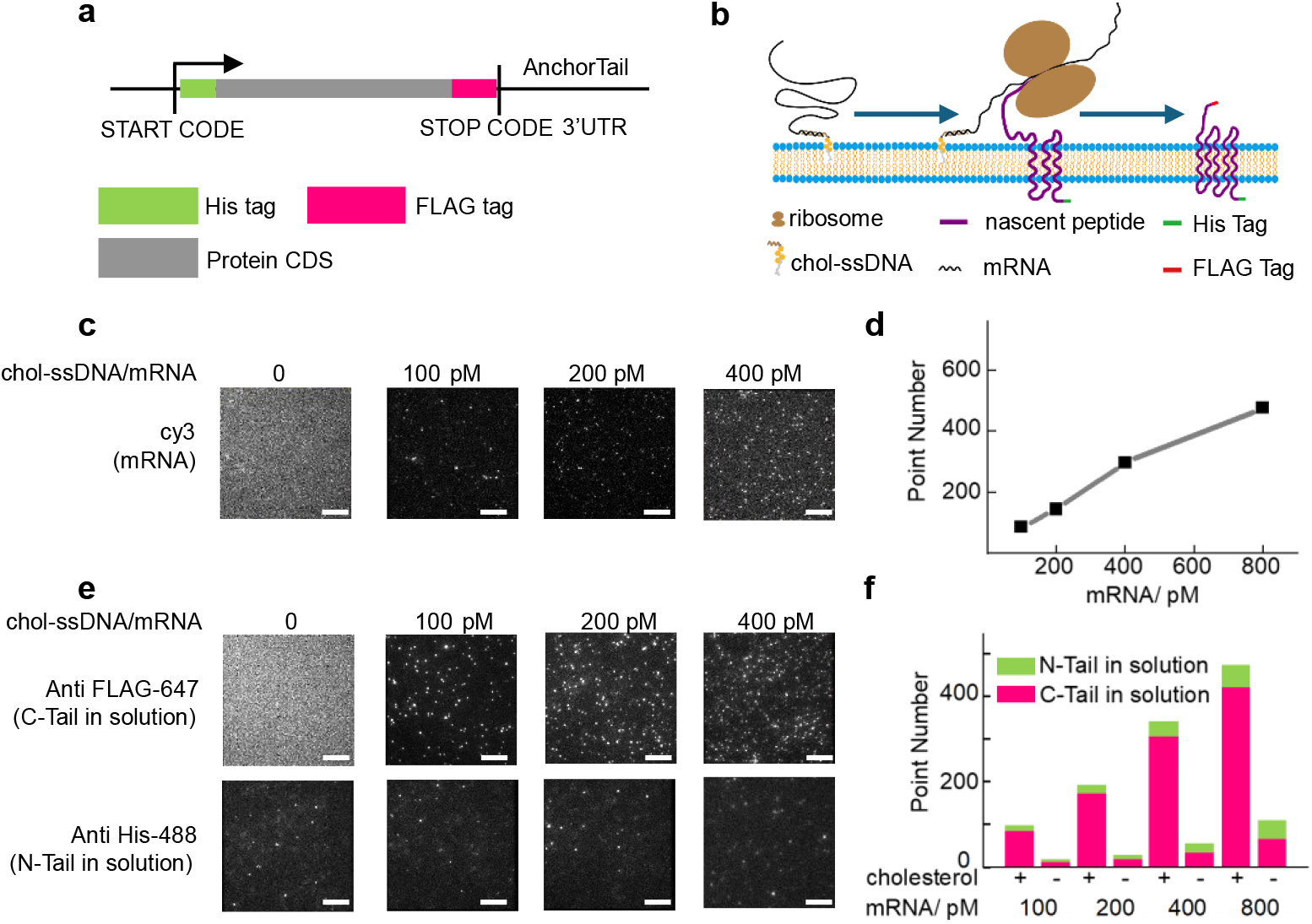
Proteins synthesized on supporting lipid bilayers via PURE. **a**, mRNA sequences with a His tag and a FLAG tag that can be used for direction detection. **b**, Schematic diagram of membrane protein synthesis by recruiting translation machinery to mRNA anchored in the supported lipid bilayer (SLB). **c**, Representative, cy3-labeled mRNA in the SLB before protein synthesis. **d**, Quantification of mRNA on the SLB. As the concentration of mRNA increased, there was more mRNA binding on the lipid bilayers. **e**, Representative fluorescent antibody signals in the SLB after F_0_ subunit a synthesis. **f**, Quantification of protein tags in the SLB. As the concentration of mRNA increases, there are more proteins in the lipid bilayers to which fluorescent antibodies can bind. The C-Tail was mainly exposed to solution. Cholesterol (+) means mRNA with cholesterol-labeled ssDNA. Cholesterol (-) refers to nucleic acids with the same structure but without cholesterol. scale bar = 10 μm.

Using the total internal reflection fluorescent (TIRF) microscopy, we observed the in vitro translation of mRNA anchored to the membrane at the single-molecule level. We prepared a glass surface coated with a phospholipid bilayer and introduced various concentrations of the synthesized mRNA for the translation and subsequent fluorescent antibody staining (detailed in Methods). This approach enabled us to determine the number and orientation of newly synthesized membrane proteins. The experimental data confirmed effective anchoring of the synthesized mRNA to the membrane (Fig. 1c). Quantitative analysis revealed that as the concentration of the cholesterol-labeled mRNA increased, so did the number of the mRNA spots on the membrane (Fig. 1d). Further quantification of single-molecule signals using different fluorescent antibodies allowed us to assess the total protein count and the orientation of the proteins on the membrane. Notably, the mRNA anchored to the membrane significantly enhanced the protein yield in the membrane. As the mRNA concentration increased, so did the number of the correctly oriented and integrated proteins. Analysis of the ratio of Anti-Flag-647 fluorescent spots to the total number of fluorescent spots showed that over 80% of the proteins were correctly integrated into the membrane at various mRNA concentrations (Fig. 1e).

### Localization enhancement of membrane proteins in synthetic cells by membrane-anchored mRNAs

After validating our strategy’s effectiveness through the SLB experiments, we extended our study to include the membrane protein synthesis within synthetic cells, employing Giant unilamellar vesicles (GUVs) to mimic cells. To demonstrate that this system is functional, we first encapsulated the cell-free translation system along with the membrane-binding mRNA inside the GUVs. We created a DNA construct that encodes atpB-mNG-AnchorTail, wherein the fluorescent protein mNeonGreen (mNG) served as a quantifiable marker. Following transcription and purification, we annealed the mRNA with the same ssDNA sequence used in the SLB experiments. We encapsulated these annealed complexes and the PURE system in GUVs via reverse emulsification at 4°C. Subsequent incubation at 37°C for three hours promoted robust membrane protein synthesis^30^. We then utilized confocal microscopy to image the post-reaction GUVs, enabling statistical analysis of the mRNA and F_0_ subunit a distribution. Imaging with Cy3-labeled mRNA confirmed substantial membrane anchoring in the presence of cholesterol (Fig 2b, c). Notably, when the mRNA was anchored to the membrane, we observed significantly enhanced localization of the ATP Synthase F_0_ subunit a-mNG fusion proteins at the membrane (Fig. 2b,c). By measuring the fluorescence intensity at the membrane compared with the fluorescence intensity within the GUVs, we observed that anchoring mRNA to the membrane led to more than a four-fold increase in the protein localization (Fig. 2d).

**Fig. 2.**
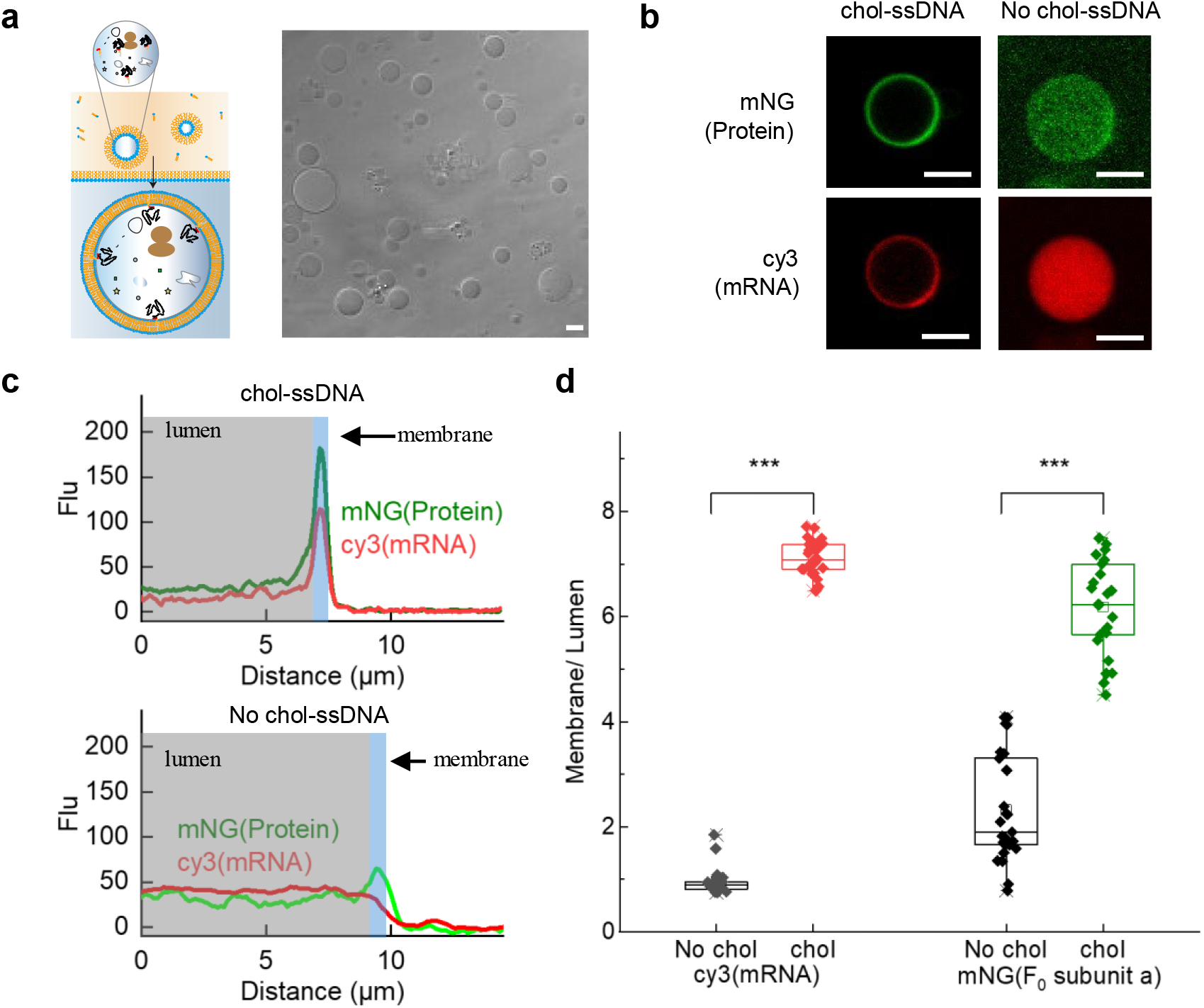
Synthesis of membrane proteins within GUVs using the PURE system. **a**, Schematic depicting the process of cell-sized GUV synthesis. **b**, Representative image of a GUV. Green fluorescence indicates the mNG-labeled F_0_ subunit a, while red fluorescence indicates the cy3-labeled mRNA. **c**, A representative fluorescence intensity profile. The grey area indicates the fluorescence intensity within the lumen, and the blue area indicates the fluorescence intensity of the membrane region. **d**, Quantification of the lumen-to-membrane fluorescence intensity ratio derived from the radial cy3 and mNG fluorescence profiles of individual GUVs. Mean value (open square), median and quartiles are shown for 25 individual GUVs for both conditions. Statistical significance was analyzed via one-way ANOVA with Tukey [HSD] post-hoc analysis, *** p < 0.001. scale bar = 10 μm.

For comparison, we also conducted the translation in a membrane-free environment and subsequently encapsulated the system into the GUVs using the same technique. The imaging demonstrated that while mRNA could be anchor to the membrane via cholesterol, almost no proteins localized to the membrane. This suggests that the F_0_ subunit a requires co-translational folding in the presence of a bilayer for effective membrane localization, a process not achievable in the GUVs created after the protein translation (Supplementary Figure 4).

The result demonstrates the necessity of our co-translational approach for efficient protein membrane integration in synthetic cell models, highlighting the potential of this method in bioengineering applications where precise protein localization is crucial.

### DNA-initiated membrane protein synthesis in synthetic cells

We have now successfully demonstrated that anchoring mRNA to the membrane significantly enhances the localization of membrane proteins within synthetic cells. In natural cells, the mRNA transcription from a DNA template initiates protein synthesis, adhering to the central dogma of molecular biology. To closely mimic this natural process, we aim to encapsulate DNA containing gene sequences, along with a cell-free expression system, inside the GUVs. This setup will allow us to assess the feasibility of synthesizing membrane proteins directly from DNA templates within synthetic cells, potentially minimizing external intervention and enhancing the cell-like functionality of our synthetic system. We incorporated cholesterol-labeled ssDNA, Cy3-labeled ssDNA and DNA templates into the PURE system and then encapsulated these complexes in the GUVs using the reverse emulsification. The cholesterol-labeled ssDNA rapidly bound to the phospholipid membrane (Supplementary Figure 5). The mRNA transcribed from the DNA templates diffuses to the membrane and spontaneously binds to the cholesterol-labeled ssDNA on the membrane, followed by translation in the vicinity of the membrane (Fig. 3a). We tested two proteins, whose templates were: atpB-mNG-AnchorTail and LacY-mNG-AnchorTail. LacY, a crucial component of the *E. coli* lactose operon with 12 transmembrane helices, facilitates lactose transport into cells using the proton motive force.

**Fig. 3.**
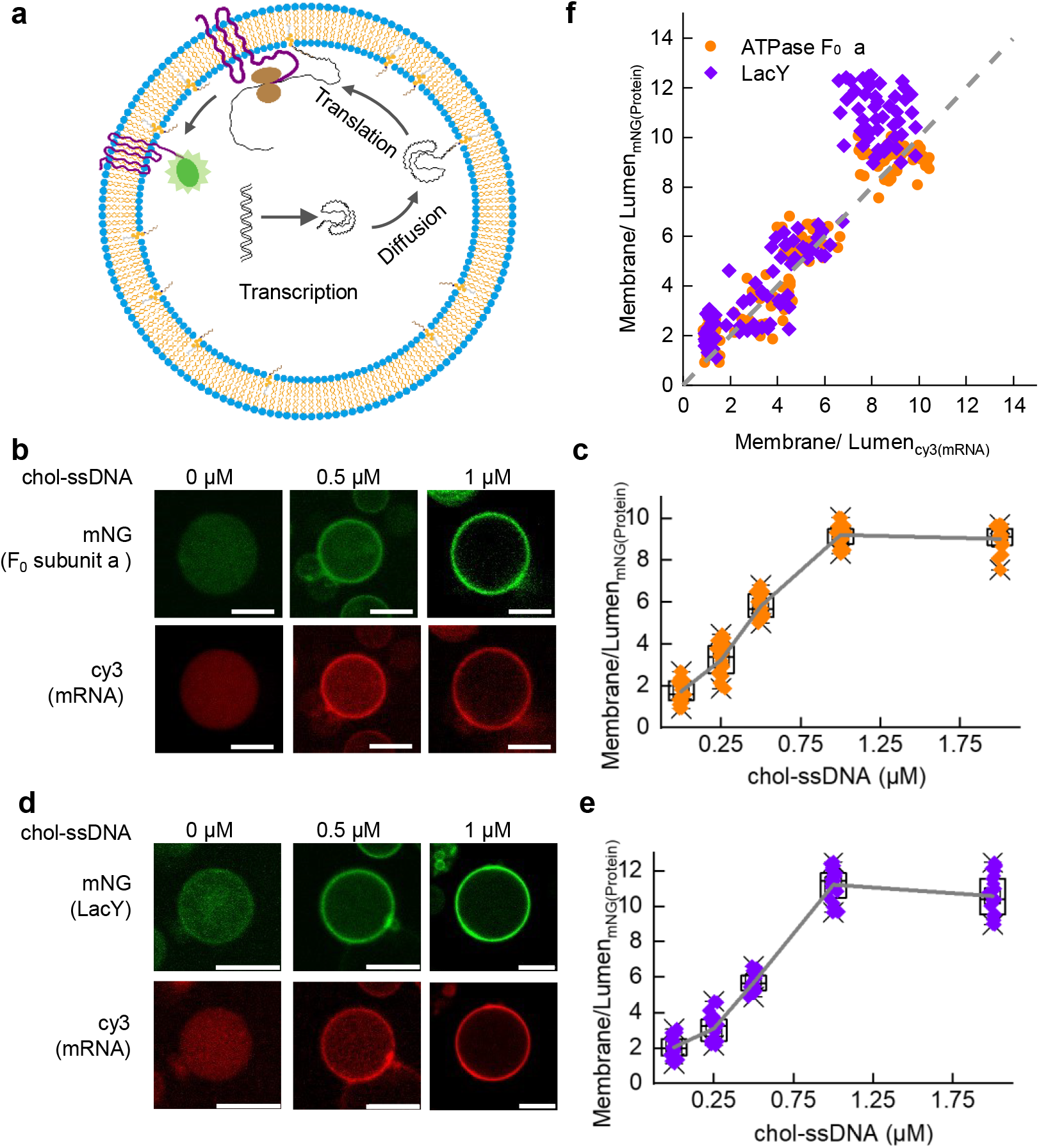
Membrane proteins synthesized in GUVs via PURE from DNA. **a**, Schematic illustrating the synthesis of membrane proteins within cell-sized GUVs. mRNA transcribed from DNA diffuses to the membrane and spontaneously binds to cholesterol-labeled ssDNA on the membrane, followed by translation in the vicinity of the membrane. **b, d**, Representative images of F_0_ subunit a **(b)** and LacY **(d)** synthesized separately in GUVs. Green fluorescence indicates the mNG-labeled protein, and red fluorescence indicates the cy3-labeled mRNA. **c, e**, Quantification of membrane/lumen fluorescence intensity ratio derived from the radial profiles of mNG fluorescence emission in individual GUVs. Mean value (open square), median and quartiles are shown for 25 individual GUVs for both conditions. **f**, Plot of the relationship between the membrane/lumen fluorescence intensity ratios of protein and mRNA in GUVs.

After incubating the GUVs at 37°C for three hours, the confocal microscopy images confirmed that the majority of the mRNA was membrane-bound, facilitated by the cholesterol-labeled ssDNA. The anchoring substantially improved the in vitro translation and the integration of the membrane proteins into the GUV membrane (Fig. 3b, d). Further analysis demonstrated a significant positive correlation between the mRNA localization and the membrane protein integration, especially, when including cholesterol (Fig. 3f). By varying the concentration of the cholesterol-labeled ssDNA, we observed enhanced mRNA membrane binding, with the membrane protein integration reaching saturation at 1 µM of cholesterol-ssDNA (Fig. 3c, e).

This approach not only demonstrated the feasibility of initiating the protein synthesis from DNA in synthetic cells but also significantly enhanced the efficiency and localization of membrane proteins. Our method lays a solid foundation for the bioengineering of complex synthetic cell systems capable of mimicking authentic cellular functions, offering a potent new tool for synthetic biology.

### Synthesis of oligomeric membrane proteins in synthetic cells

Encouraged by the results above, we decided to synthesize small oligomeric membrane proteins. We used EmrE as a model protein, which is a multidrug transporter in *E. coli*, consisting of 110 amino acids that constitute its four transmembrane domains. The EmrE proteins form antiparallel dimers in the membrane to be functional (Supplementary Figure 6)^31–33^. Due to its small size, dual topology and high hydrophobicity, EmrE has been reported to appear in the insoluble fraction when synthesized using in vitro translation. Soga and colleagues achieved a substantial increase in the integration rate by reducing the size of their synthetic cells, effectively increasing the phospholipid concentration around the translation machinery^17^. Here, anchoring the mRNA to the membrane offers a versatile solution to produce the membrane proteins in synthetic cells without limiting their sizes.

We employed the transmembrane dye FlAsH, which binds covalently to the tetracysteine tag (TC-Tag) and generates fluorescence, to track the EmrE protein localization and ratio during the in vitro synthesis. We refrained from using fluorescent proteins like GFP because the large fluorescent protein might disturb the dimerization and the topological inversion of EmrE. We synthesized the mRNA encoding EmrE with a TC-tag and an AnchorTail sequence. The translation experiments on SLBs showed that this approach supports the synthesis and dimerization of EmrE. Moreover, enhancing the concentration of the membrane-bound mRNA directly led to more EmrE production (Fig. 4a). We employed single-molecule photobleaching step analysis to quantify the number of the subunits within the protein complexes. The subunit counting relies on identifying individual bleach steps of fluorescently tagged proteins-of-interest (Fig. 4b). The statistical analysis revealed that the proportion of two-step photobleaching events was higher than expected from random formation processes (Fig. 4c), suggesting the formation a dimeric structure.

**Fig. 4.**
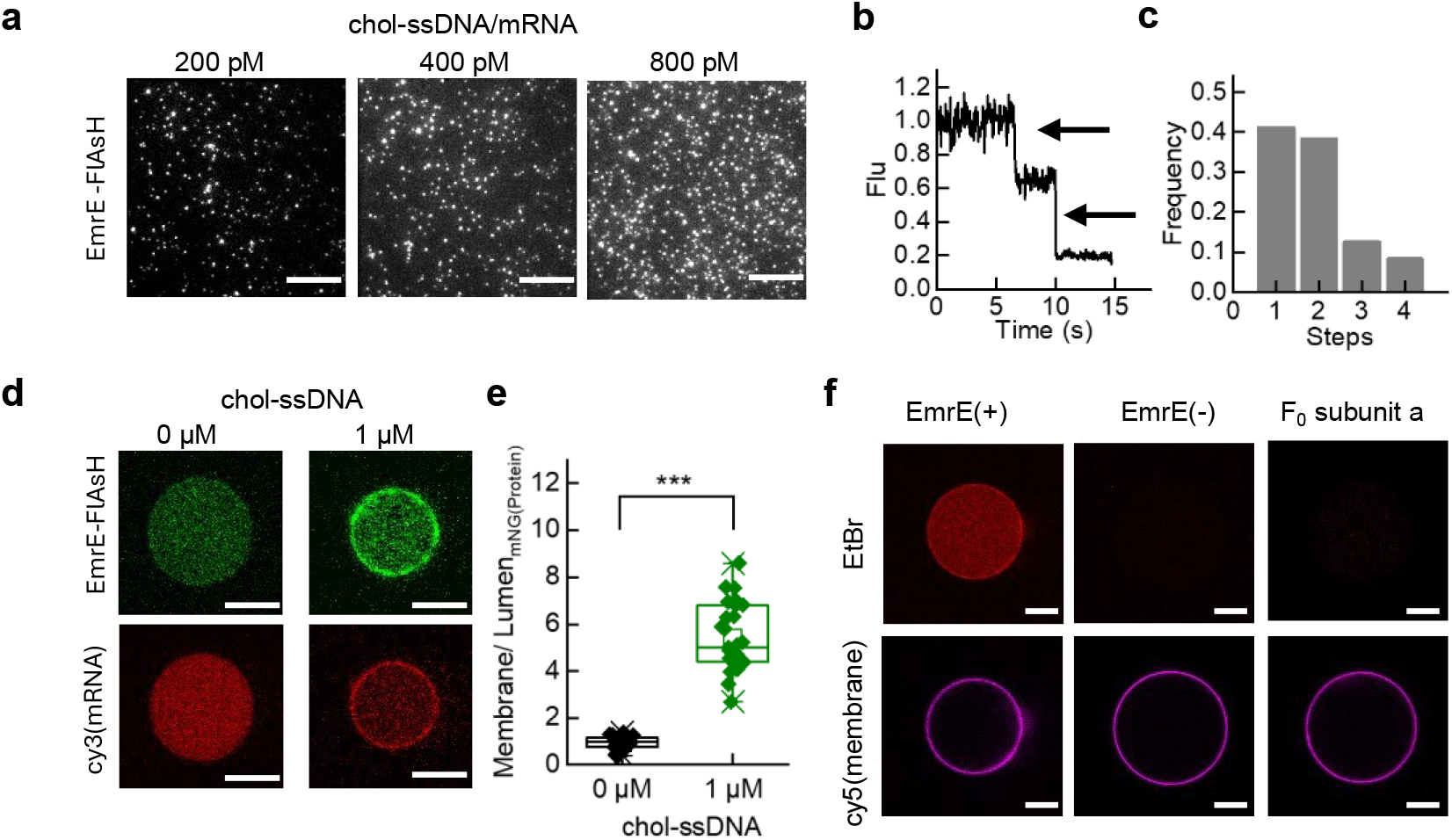
Synthesis of oligomeric membrane proteins in large synthetic cells. **a**, Representative TIRF image of EmrE-FlAsH signals in a SLB after protein synthesis. **b**, Representative intensity−time trace of the photobleaching steps observed for EmrE in the SLB. The two photobleaching steps are denoted by arrows. **c**, Histogram of the number of photobleaching steps from the intensity−time trace, n = 256 spots. **d**, Typical GUV image. Green fluorescence indicates the FlAsH-labeled EmrE, and red fluorescence indicates the cy3-labeled mRNA. **e**, Quantification of the FlAsH fluorescence intensity from a single GUV. Mean value (open square), median and quartiles are shown for 26 individual GUVs for both conditions. Statistical significance was analyzed via one-way ANOVA with Tukey [HSD] post-hoc analysis, *** p < 0.001. **f**, Representative microscopy images of EmrE transporting EtBr into a GUV.

We performed the protein synthesis using the same DNA template as above and stained the newly synthesized proteins for quantitative analysis using FlAsH. We specifically chose the synthetic cells exceeding 10 µm for the statistics. Previous studies have indicated that the integration rate of EmrE into membranes in synthetic cells of this size is less than 1%^17^. As illustrated by our data (Fig. 4d), without the use of the cholesterol-labeled ssDNA, the fluorescence intensity on the membrane does not significantly differ from the fluorescence intensity in the lumen. However, with the assistance of the cholesterol-labeled ssDNA, the mRNA carrying the AnchorTail sequence is recruited to the membrane, significantly enhancing the proportion of EmrE on the membrane (Fig. 4 d, e).

We performed a functional test of the protein using ethidium bromide (EtBr) as a model substrate, which fluoresces upon entering the GUVs and binding to internal nucleic acids. After synthesizing EmrE, we introduced EtBr into the external solution of the GUVs and established a pH gradient to facilitate its transport. EmrE’s functionality was confirmed by its ability to transport EtBr into the GUVs, evidenced by the significantly brighter fluorescence in the GUVs expressing EmrE compared to the controls that expressing the F_0_ subunit a (Fig. 4f). The results not only demonstrated the successful integration and functional activity of EmrE within the synthetic cells but also confirmed its existence as a functional, antiparallel homodimer. Our methodology therefore offers a robust platform for the bioengineering of functional oligomeric membrane proteins in synthetic cells.

## Discussion

This study introduces a novel method that spatially confines mRNA to the membrane by incorporation of a specific sequence into its 3’ UTR, thereby enabling the translation processes to occur close to the membrane even within synthetic cells tens of micrometers in diameter that are composed of GUVs encapsulating the membrane protein expression systems and genetic constructs. This membrane-proximity facilitates direct interaction between the proteins and the phospholipid membrane and hence co-translational folding. Our results demonstrated that cholesterol-labeled single-stranded DNA can effectively bind to the non-coding regions of mRNA, substantially increasing the concentration of mRNA at the membrane interface available for translation. This method has demonstrated a six-to ten-fold increase in the membrane binding efficiency of newly synthesized proteins compared with that under the control conditions. Compared with the methods that increase the liposome concentration or reduce the protocell volume to promote the membrane protein integration, our method is more universally applicable. By editing the 3’ UTR sequences, the integration of specific membrane proteins in the membrane is facilitated independently of the cell size.

Importantly, our method does not require modification to the coding sequences (CDS) and is compatible with standard commercial in vitro transcription-translation systems (IVTTs). The utilization of the 3’ UTR for molecular recognition makes our technology highly programmable and versatile, enabling region-specific protein expression within synthetic cells. Our approach permits the translation of membrane proteins near the membrane, while water-soluble proteins are translated in the cytoplasmic equivalent regions.

Although our experiments leveraged cholesterol-labeled ssDNA that annealed with the 3’ UTR, this method could in principle be adapted to use any DNA, protein or other molecules that can specifically bind to the 3’ UTR. This adaptability highlights the significant potential and expandability of our approach, which is poised to revolutionize the field of synthetic biology by providing a robust platform for the targeted synthesis and functional integration of membrane proteins in artificial cell systems.

## Method

### Material

In this study, all phospholipids were purchased from Avanti Polar Lipids. The in vitro transcription kit was obtained from M0251S (New England Biolabs, NEB). The PURE system was sourced from PURE*frex*®2.1 (GeneFrontier). Unless otherwise noted, all sequences were synthesized by Sangon Biotech, and all chemicals were procured from Sigma-Aldrich.

### Preparation of Template DNA and mRNA

Plasmid DNA (pET28-atpB-mNG and pET28-LacY-mNG) containing gene coding for ATP Synthase F_0_ subunit a-mNG and LacY-mNG from *E. coli* MG1655 was synthesized by BGI Company. These vectors include a T7 promoter and ribosome binding site. The template DNA was amplified by PCR, adding Tag and AnchorTail, with the sequences of all primers listed in Supplementary Information (SI) Table 1. The PCR products were verified by sequencing. mRNA was synthesized in vitro using a standard T7 RNA polymerase protocol. For purification, 100 μL IVT solution was mixed with 900 μL TRIS, extracted with 200 μL chloroform, and precipitated with isopropyl alcohol and sodium acetate. The RNA pellet was washed with 75% ethanol, resuspended in RNase-free water, and stored at -80°C.

### Cell-Free Protein Synthesis in Supported Lipid Bilayer and TIRF Microscopy

Small unilamellar vesicles (SUVs) composed of POPC were prepared at 2 mg/mL using the thin film hydration method. POPC in chloroform was evaporated under nitrogen, dried under vacuum, and hydrated with 5 mL milli-Q PBS at 37°C, followed by sonication until clear. Cleaned microscope cover glass was used to form supported lipid bilayers by depositing and incubating 80 μL of SUV solution for six hours, followed by rinsing with PBS.

A custom-built objective-TIRF microscope was utilized for single-molecule imaging at wavelengths specific for different fluorophores. For cell-free protein synthesis on SLBs, annealed mRNA was diluted and added to PMMA wells with SLB. After incubation at room temperature, the 2X PURE*frex* system was added and the mixture incubated at 37°C. Post-synthesis, samples were treated with RNase and rinsed with PBS. Proteins on SLBs were stained with Anti-His-488, Anti-Flag-647, or FlAsH-EDT2 and imaged using TIRF microscopy to quantify protein number and orientation.

### Cell-Free Protein Synthesis in GUV

For synthesis within GUVs, the w/o emulsion-transfer method was employed. Briefly, 20 μL of the PURE system supplemented with template DNA, 200 mM sucrose, 0.8 U/ μL of RNase inhibitor, 50 nM cy3-labeled ssDNA, and cholesterol (±) labeled ssDNA were added to 600 μL of mineral oil containing 0.5 mg/mL POPC. The mixture was vortexed for 30 seconds to form a w/o emulsion and then balanced on ice for 10 minutes. An aliquot of 600 μL of this solution was gently placed on top of 200 μL of the outer solution (see composition below) and centrifuged at 2,000g for 10 minutes at 4°C. The pelleted GUVs were collected through an opening at the bottom of the tube and suspended in fresh outer solution. Protein synthesis inside the GUVs was conducted at 37°C for 3 hours. The outer solution contained the low-molecular-weight components of the PURE system as well as 200 mM glucose to balance osmotic pressure but did not include ribosomes or proteins needed for transcription and translation processes. To detect the influx of EtBr, the external solution of the GUVs was replaced with a buffer that was identical to the dilution buffer but contained 10 μM EtBr at a pH of 8.1.

### Confocal Microscopy

Images were captured using an Olympus FV3000 confocal microscope. Fiji software was used for fluorescence image analysis, with background adjustments made using predefined ROIs near each GUV.

## Statistical Analysis

Data were analyzed using one-way ANOVA with Tukey’s post-hoc test and two-tailed, two-sample Student’s t-test, performed using Python version 3.10. Statistical significance was assumed for p-values < 0.001 unless stated otherwise.

## Author contributions

Y.L., M.L and H.F. conceived of and designed the project. H.F., L.M., C,X., J.L., Y.S., and Y,T. performed and analyzed experiments. Y.L., M.L and H.F. wrote the manuscript. S.H., Y.Q., M.F., H.Z., C.L., and F.Y. provided technical support for liposome preparation and microscope imaging.

## Acknowledgements

This work was supported by the Strategic Priority Research Program of the Chinese Academy of Sciences (XDB0480000); the National Key Research and Development Programme of China (2019YFA0709304); and National Natural Science Foundation of China (T2221001, 32171228, 32471278)

## References

1. Yıldırım, M. A., Goh, K.-I., Cusick, M. E., Barabási, A.-L. & Vidal, M. Drug— target network. Nat Biotechnol 25, 1119–1126 (2007).

2. Kohyama, S., Merino-Salomón, A. & Schwille, P. In vitro assembly, positioning and contraction of a division ring in minimal cells. Nat Commun 13, 6098 (2022).

3. Lee, K. Y. et al. Photosynthetic artificial organelles sustain and control ATP-dependent reactions in a protocellular system. Nat Biotechnol 36, 530–535 (2018).

4. Berhanu, S., Ueda, T. & Kuruma, Y. Artificial photosynthetic cell producing energy for protein synthesis. Nat Commun 10, 1325 (2019).

5. Grisshammer, R. Understanding recombinant expression of membrane proteins. Current Opinion in Biotechnology 17, 337–340 (2006).

6. Cymer, F., Von Heijne, G. & White, S. H. Mechanisms of Integral Membrane Protein Insertion and Folding. Journal of Molecular Biology 427, 999–1022 (2015).

7. Wang, S. et al. The molecular mechanism of cotranslational membrane protein recognition and targeting by SecA. Nature Structural & Molecular Biology 26, 919–929 (2019).

8. Ismail, N., Hedman, R., Lindén, M. & von Heijne, G. Charge-driven dynamics of nascent-chain movement through the SecYEG translocon. Nature Structural & Molecular Biology 22, 145–149 (2015).

9. Frauenfeld, J. et al. Cryo-EM structure of the ribosome–SecYE complex in the membrane environment. Nature Structural & Molecular Biology 18, 614–621 (2011).

10. Steinberg, R., Knüpffer, L., Origi, A., Asti, R. & Koch, H.-G. Co-translational protein targeting in bacteria. FEMS Microbiology Letters 365, (2018).

11. Volkov, I. L. et al. Spatiotemporal kinetics of the SRP pathway in live E. coli cells. Proceedings of the National Academy of Sciences 119, e2204038119 (2022).

12. Schibich, D. et al. Global profiling of SRP interaction with nascent polypeptides. Nature 536, 219–223 (2016).

13. Wimley, W. C. & White, S. H. Experimentally determined hydrophobicity scale for proteins at membrane interfaces. Nat Struct Mol Biol 3, 842–848 (1996).

14. Ulmschneider, J. P., Smith, J. C., White, S. H. & Ulmschneider, M. B. In Silico Partitioning and Transmembrane Insertion of Hydrophobic Peptides under Equilibrium Conditions. J. Am. Chem. Soc. 133, 15487–15495 (2011).

15. Almeida, P. F., Ladokhin, A. S. & White, S. H. Hydrogen-bond energetics drive helix formation in membrane interfaces. Biochimica et Biophysica Acta (BBA) - Biomembranes 1818, 178–182 (2012).

16. MacKenzie, K. R. Folding and Stability of α-Helical Integral Membrane Proteins. Chem. Rev. 106, 1931–1977 (2006).

17. Soga, H. et al. In Vitro Membrane Protein Synthesis Inside Cell-Sized Vesicles Reveals the Dependence of Membrane Protein Integration on Vesicle Volume. ACS Synth. Biol. 3, 372–379 (2014).

18. Ando, M., Schikula, S., Sasaki, Y. & Akiyoshi, K. Proteoliposome Engineering with Cell-Free Membrane Protein Synthesis: Control of Membrane Protein Sorting into Liposomes by Chaperoning Systems. Advanced Science 5, 1800524 (2018).

19. Eaglesfield, R., Madsen, M. A., Sanyal, S., Reboud, J. & Amtmann, A. Cotranslational recruitment of ribosomes in protocells recreates a translocon-independent mechanism of proteorhodopsin biogenesis. iScience 24, 102429 (2021).

20. Lawrence, J. Intracellular localization of messenger RNAs for cytoskeletal proteins. Cell 45, 407–415 (1986).

21. Das, S., Vera, M., Gandin, V., Singer, R. H. & Tutucci, E. Intracellular mRNA transport and localized translation. Nat Rev Mol Cell Biol 22, 483–504 (2021).

22. Korkmazhan, E., Teimouri, H., Peterman, N. & Levine, E. Dynamics of translation can determine the spatial organization of membrane-bound proteins and their mRNA. Proc. Natl. Acad. Sci. U.S.A. 114, 13424–13429 (2017).

23. Xia, C., Fan, J., Emanuel, G., Hao, J. & Zhuang, X. Spatial transcriptome profiling by MERFISH reveals subcellular RNA compartmentalization and cell cycle-dependent gene expression. Proc. Natl. Acad. Sci. U.S.A. 116, 19490–19499 (2019).

24. Mili, S., Moissoglu, K. & Macara, I. G. Genome-wide screen reveals APC-associated RNAs enriched in cell protrusions. Nature 453, 115–119 (2008).

25. Reid, D. W. & Nicchitta, C. V. Diversity and selectivity in mRNA translation on the endoplasmic reticulum. Nat Rev Mol Cell Biol 16, 221–231 (2015).

26. Villanueva, E. et al. System-wide analysis of RNA and protein subcellular localization dynamics. Nat Methods 21, 60–71 (2024).

27. Cui, X. A., Zhang, H. & Palazzo, A. F. p180 Promotes the Ribosome-Independent Localization of a Subset of mRNA to the Endoplasmic Reticulum. PLoS Biol 10, e1001336 (2012).

28. Sarmah, P. et al. mRNA targeting eliminates the need for the signal recognition particle during membrane protein insertion in bacteria. Cell Reports 42, 112140 (2023).

29. Nevo-Dinur, K., Nussbaum-Shochat, A., Ben-Yehuda, S. & Amster-Choder, O. Translation-Independent Localization of mRNA in E. coli. Science 331, 1081–1084 (2011).

30. Fujii, S. et al. Liposome display for in vitro selection and evolution of membrane proteins. Nat Protoc 9, 1578–1591 (2014).

31. Fluman, N., Tobiasson, V. & Von Heijne, G. Stable membrane orientations of small dual-topology membrane proteins. Proc. Natl. Acad. Sci. U.S.A. 114, 7987–7992 (2017).

32. Seurig, M., Ek, M., Von Heijne, G. & Fluman, N. Dynamic membrane topology in an unassembled membrane protein. Nat Chem Biol 15, 945–948 (2019).

33. Woodall, N. B., Yin, Y. & Bowie, J. U. Dual-topology insertion of a dual-topology membrane protein. Nat Commun 6, 8099 (2015).

