## Supplementary Figures and Table for "*In situ* Membrane Protein Expression by Efficient Recruitment of mRNA to the Membranes of Synthetic Cells"

Supp. Fig. 1: The topology of His and FLAG labeled ATP Synthase $F_0$ subunit a
Supp. Fig. 2: The topology of membrane-binding mRNA.
Supp. Fig. 3: Examples of radial profile fluorescence analysis.
Supp. Fig. 4: Membrane recruitment and insertion of ATPB and LacY does not occur post-translationally.
Supp. Fig. 5: Cholesterol-labeled ssDNA associated with GUVs can quickly adhere to the membrane

### Supplementary Table

Supp. Tab. 1: All sequences used in this study.

### Supplementary Figures

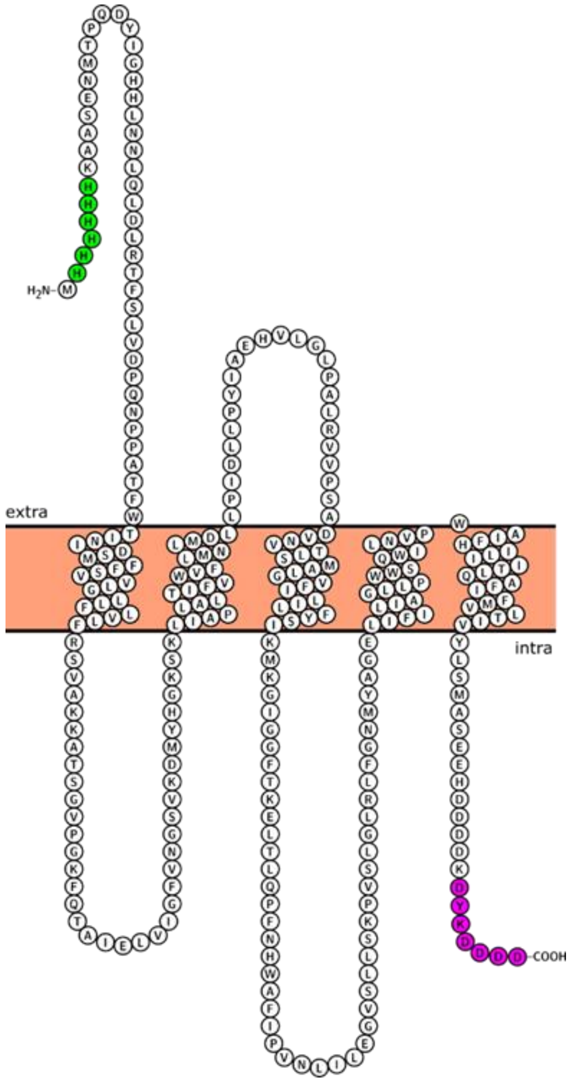


Supplementary Figures 1.The topology of His(green) and FLAG(magenta) labeled ATP Synthase $F_0$ subunit a. The F0 subunit a contains five transmembrane helices, and proper binding to the membrane results in the N- and C-termini being exposed on opposite side. When it is correctly folded, the C-termini are on the same side as the translational elements.

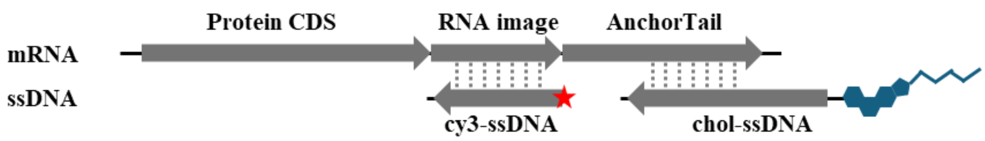


Supplementary Figures 2. The topology of membrane-binding mRNA. The arrow shows the direction from the 5' to the 3' end."

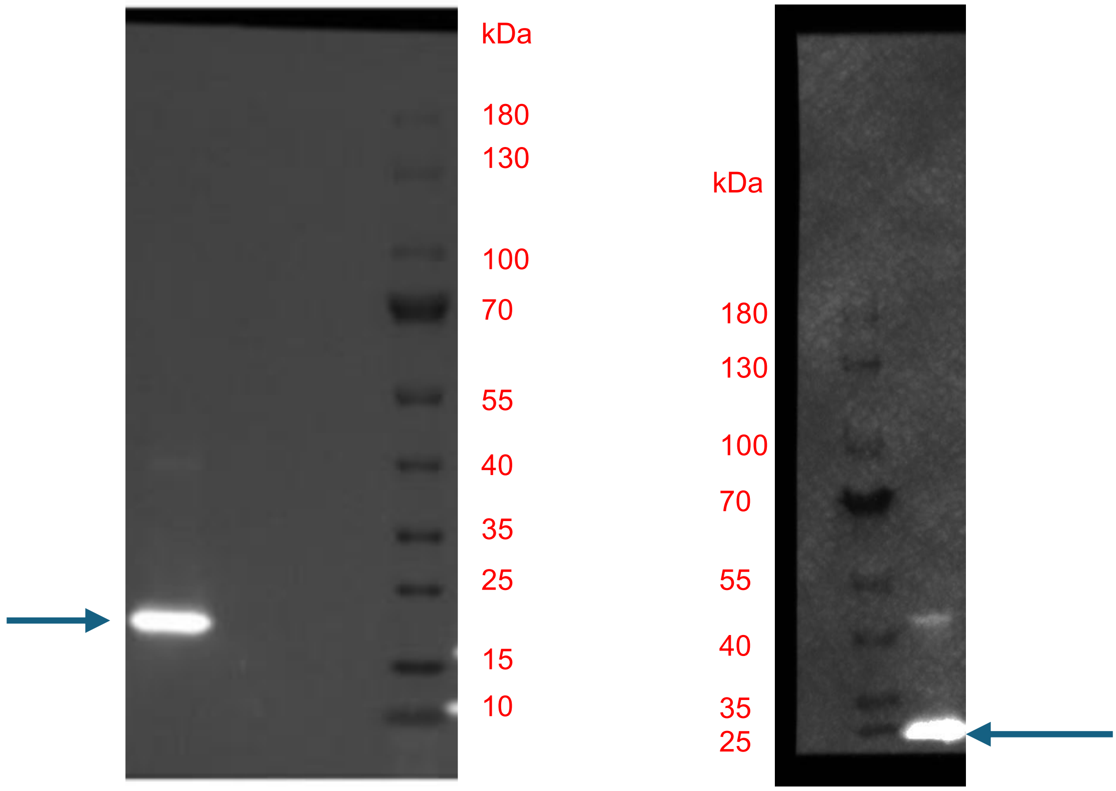


Supplementary Figures 3.Western blot analysis of His-atpB-FLAG synthesized by cell-free protein synthesis in the bulk system. Immunoblotting was performed using AntiHis-647(left) or AntiFlag-488(right) antibodies.

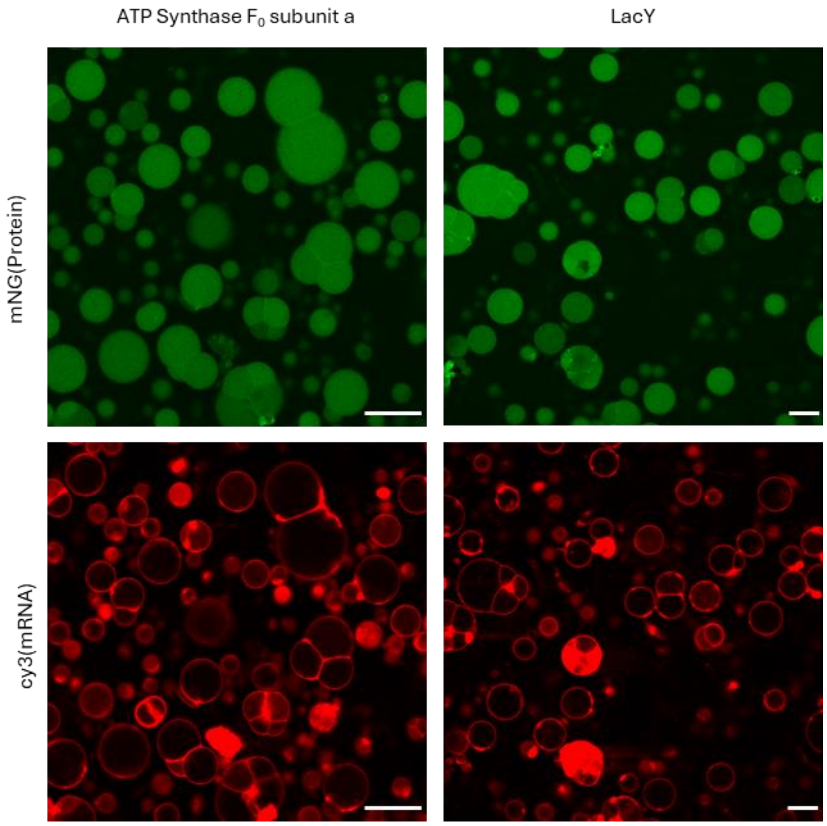


Supplementary Figures 4. Membrane recruitment and insertion of ATP Synthase $F_0$ subunit a and LacY does not occur post-translationally. Scale bar=30 µm

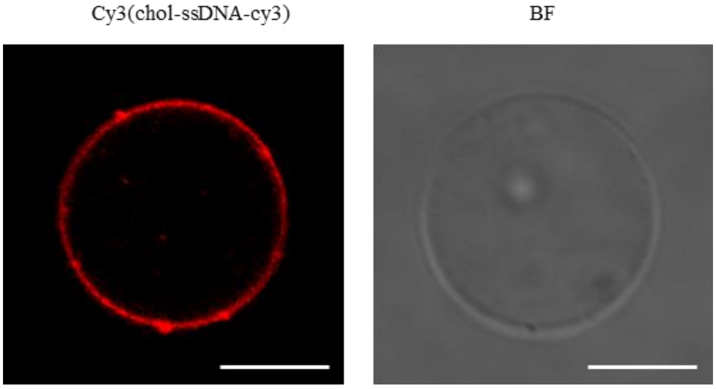


Supplementary Figures 5. Cholesterol-labeled ssDNA associated with GUVs can quickly adhere to the membrane. BF means bright-field. Scale bar =10 μm.

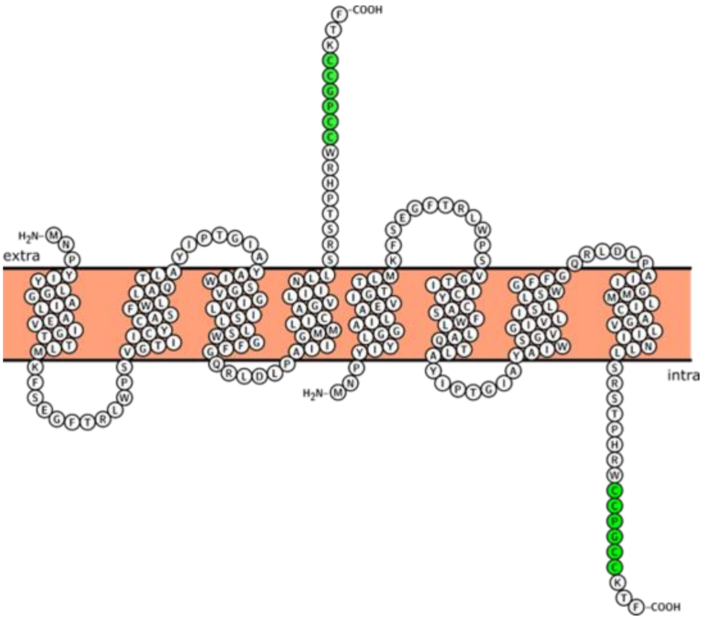


Supplementary Figures 6.The topology of tetracysteine tag(green) labeled EmrE dimer.

| | Sequence |
|---|---|
| AnchorTail | (Protein CDS)-CCTTGGAAATGGTTCGGGCACTACACCGTGGTGGAGCAATGTCATCGATAGGGTCACTCGTCTTGACGGCATGTGGAAGGTAA |
| chol-ssDNA | (chol)TTTTTTTTTTCCTTCCACATGCCGTCAAGACGAGTGACCC |
| chol-ssDNA-cy3 | (chol)TTTTTTTTTTCCTTCCACATGCCGTCAAGACGAGTGACCC(cy3) |
| cy3-ssDNA | (cy3)ATTGCTCCACCACGGTGTAG |

Supplementary Table 1: All sequences used in this study.
